# Saccadic adaptation in the presence of artificial central scotomas

**DOI:** 10.1101/719450

**Authors:** Youngmin Song, Lydia Ouchene, Aarlenne Z. Khan

## Abstract

Saccadic adaptation can occur over a short period of time through a constant adjustment of the saccade target during the saccade, resulting in saccadic re-referencing which directs the saccade to a location different from the target that elicited the saccade. Saccade re-referencing could be used to help patients with age-related macular degeneration (AMD) to optimally use their residual visual function. However, it remains unknown whether saccade adaptation can take place in the presence of central scotomas (i.e., without central vision).

We tested participants in two experiments in a conventional double-step paradigm with a central gaze-contingent artificial scotoma. Experiment 1 (*N* = 12) comprised a backward adaptation paradigm with a visible and an invisible 3° diameter scotomas. Experiment 2 (*N* = 13) comprised a forward adaptation paradigm with invisible 2° and 4° diameter scotomas.

In Experiment 1, we observed significant adaptation in both the visible and invisible scotoma conditions comparable to the control condition with no scotoma. This was the case even when the saccade landed such that the target was occluded by the scotoma. We observed that adaptation occurred based on peripheral viewing of the stepped target during the deceleration period.

In Experiment 2, we found that both scotoma conditions showed adaptation again comparable to the control condition with no scotoma. We conclude that saccadic adaptation can occur with central scotomas, showing that it does not require central vision and is driven primarily by peripheral retinal error.

## Introduction

Saccades continue to be accurate even as one ages and extraocular muscles weaken because the brain monitors and adjusts the accuracy of saccades (Herman et al., 2013). This is known as saccadic adaptation. For example, saccades that undershoot or overshoot a target are adjusted over a short period of time so that subsequent saccades will have an amplitude closer to the target’s distance. This is easily demonstrable in a lab setting using a double-step paradigm (McLaughlin, 1967) in which a target’s position is repeatedly shifted during the saccade. Gradually, saccades land closer to the shifted target position.

Saccadic adaptation could be used for rehabilitation of patients with age-related macular degeneration (AMD). In AMD, deterioration of the macula impedes visual acuity by affecting central vision (Cacho et al., 2010). Due to this, patients need to use their intact peripheral vision to access visual information. They can be trained or can spontaneously learn to consistently use one or more peripheral regions, known as preferred retinal loci (PRL) (Cheung & Legge, 2005; Chung, 2013; Crossland et al., 2005; Fletcher & Schuchard, 1997). Saccade re-referencing in which eye movements direct the PRL instead of the fovea to the object would greatly improve visual abilities (Cheung & Legge, 2005; Nilsson et al., 1998; Sunness et al., 1996; Walsh & Liu, 2014; White & Bedell, 1990), speeding up visual discrimination in the periphery. However, directing the PRL to the object instead has shown to be extremely difficult and if successful, can take years (Cheung & Legge, 2005; Krauzlis et al., 2017). One possibility to achieve saccade re-referencing is through training using saccadic adaptation. Here, we investigate whether saccadic adaptation can occur in the presence of a central scotoma, and if so to what extent.

Despite the multitude of studies on saccadic adaptation, the nature of the error signal that drives it has not been fully resolved. If central vision is necessary for saccadic adaptation, then it is unlikely to occur with a central scotoma. Many studies have indirectly shown that central vision is unnecessary for adaptation. For instance, some have shown that similar amounts of adaptation occurred even when the task was modified to elicit very few corrective saccades (Noto & Robinson, 2001; Wallman & Fuchs, 1998). Thus, feedback based on central vision after the corrective saccade is not necessary for adaptation. Similarly, other studies have suggested that adaptation occurs in response to a *peripheral* retinal error after the first saccade (Noto & Robinson, 2001; Wallman & Fuchs, 1998) or a difference between the post-saccadic retinal image and predicted image, again based entirely on *peripheral vision* of the desired target (Bahcall & Kowler, 2000). For example, Bahcall and Kowler (2000) demonstrated backward saccadic adaptation during a task in which participants were instructed to saccade partway to a target (75% of the distance from initial fixation point), which could not be based on retinal error. Furthermore, recent evidence has shown that even intra-saccadic visual feedback received mid-flight during a saccade is sufficient to result in saccadic adaptation (Panouillères et al., 2016; Panouillères et al., 2013). These findings support that peripheral visual information is used for adaptation rather than central. However, notably, it has not yet been demonstrated that occlusion of central vision does not impact adaptation in any way. It may be that central vision (for example after the corrective saccade or once adaptation has occurred) plays a role in adaptation, such as determining when to stop adapting. Occlusion of the target after the corrective saccade might be interpreted as a change in the external visual scene, which might also impact saccadic adaptation.

It is unclear whether there are limits to the eccentricity of peripheral visual information that can drive adaptation. If so, different sized scotomas may have different influences on adaptation. Robinson et al. (2003) tested saccadic adaptation in monkeys and showed that adaptation was most consistent for target shifts of 20 to 60% of the target eccentricity, with a decrease in adaptation for greater eccentricities (although not for forward adaptation), as well as inconsistent adaptation for smaller target shifts (<20%). However, this has not been tested in humans, who show quicker adaptation as well as stronger effects compared to monkeys (Albano & King, 1989; Deubel et al., 1986; Straube et al., 1997). Also, it should be noted that the number of adaptation trials was extensive (400 to 2,800). With human participants and fewer trials, it is uncertain whether larger scotomas would result in the shifted target being occluded sooner during the saccade and thus reduce the amount of adaptation. We therefore tested if changing the size of the scotoma influences adaptation.

While adaptation has been shown to occur in both backward and forward target shifts, there are many differences between backward and forward adaptation. For one, forward adaptation is less efficient, takes longer, and results in less gain change compared to backward adaptation (Ethier et al., 2008; Hernandez et al., 2008; Panouillères et al., 2009; Straube & Deubel, 1995). But more importantly, there is both behavioural and neurological evidence that they have different underlying neuronal mechanisms (Pélisson et al., 2010). A popular model is that while backward adaptation is caused by a decrease in saccade gain, forward adaptation relies on a remapping mechanism (Ethier et al., 2008; Hernandez et al., 2008; Semmlow et al., 1989). Neurological evidence also suggests a difference in mechanism such as Purkinje cells in the cerebellum firing differently in forward and backward adaptation (Catz et al., 2008) and forward adaptation being more affected by cerebral lesions than backward (Golla et al., 2008). Therefore, we tested both paradigms with central scotomas to determine if there are any differences in the amount of adaptation.

We also tested whether saccadic adaptation is impacted by the visibility of the scotoma. For example, a visible scotoma provides continuous feedback of the eye position during the adaptation task and may negatively impact adaptation since it provides more accurate information about the target position and shifts relative to eye position.

In summary, we investigated whether adaptation can occur in response to only peripherally viewed targets in the presence of an artificial central scotoma. In Experiment 1, we used a backward adaptation paradigm and varied the visibility of the scotoma (visible and invisible). In Experiment 2, we used a forward adaptation paradigm and tested invisible scotomas of two different sizes (2° and 4°). We found that in both experiments with central scotomas, saccadic adaptation occurred to a degree similar to those in the control conditions.

## Experiment 1

### Methods

#### Participants

Twelve participants took part in this study (three male, age range: 19-40, *M* = 22.92, *SD* = 5.52, including two authors AK and LO). All participants had normal or corrected-to-normal vision and no known neurological impairments. All gave written informed consent to participate in the experiment. All procedures were pre-approved by the Health Research Ethics Committee at the University of Montreal. (16-129-CERES-D).

#### Apparatus

Participants sat in a dark room facing a VIEWPixx LCD monitor (VPixx Technologies, Montreal, QC, 120 Hz), its center aligned horizontally with the participant’s mid-sagittal plane and vertically at eye level. The screen dimensions were 52.1 cm by 29.2 cm. The screen was 62 cm from the participant’s eyes. The participants’ heads were immobilized via a chin and forehead support placed at the edge of the table on which the monitor was located. Eye-movements were recorded using an infrared-emitting video-based eye tracker (EyeLink 1000 Plus, SR Research, Mississauga, ON, Canada). The backlight on the screen was set to a very low setting to ensure that the monitor frame was not easily visible (ViewPixx, back light setting: 5). The position of the right eye was recorded at 1000 Hz using the Eyelink 1000 video-based eye tracker (SR Research).

The scotoma was centered on the participant’s foveal vision based on a 2-step calibration process at the beginning of each experimental session. First, the standard nine-point Eyelink calibration/validation procedure was performed tracking the participant’s right eye. A second 30-point calibration was then performed to calculate horizontal and vertical correction values to precisely align the scotoma on the fovea of the participant. Participants were asked to look at each of the 15 fixation discs (black on white background, 0.25° diameter spanning 4/5^th^ of the screen) which were presented in random order and press a button when they were fixating accurately. A custom code mapped the eye positions to the fixation disc locations using a polynomial regression with 6 parameters. These parameters were then used to adjust eye position for scotoma presentation. The standard calibration resulted in a mean error of 2.21° in absolute distance (distance between recorded position and the actual gaze position) while the second calibration used for the artificial scotoma position reduced this error to 0.53° in absolute distance.

In terms of timing, the minimum delay between the position of the eye and the scotoma was approximately 3 ms and the maximum was 8.3 ms. The end-to-end sample delay for eye recording at 1000 Hz was 1.95 ms (SR Research, Kanata, Canada). In addition, there was approximately 1 ms between the time at which the eye position was determined and the rendering of the scotoma. As the screen refresh rate (120 Hz/8.3 ms) was slower than all eye-tracking and artificial-scotoma-related delays, these delays necessarily went unnoticed by the participants. Nonetheless, to ensure the precise tracking of our participants’ eyes, we ordered the computer to use the first sample from the eye-tracker following each new frame drawing to position the scotoma. Such an approach allowed us to ensure that the scotoma would be updated for every single frame. Considering that our eye-tracker was recording at 1000 Hz, we would expect 8 (or 9 in certain cases) eye tracking samples in-between frames. Finally, we wished to prevent the scotoma from following the eye during blinks (Aguilar & Castet, 2011); to do so, we froze the scotoma in place and blurred the screen whenever the velocity in the vertical direction exceeded 900°/sec or when eye movement was not detected.

#### Procedure

Stimuli used are shown in Figure 1. A white oval fixation stimulus was used instead of a small dot or cross to ensure that participants would be able to fixate even in the presence of the scotoma. It was located 4.9° left of the center horizontally, at the center of the screen vertically, and was 0.9° by 4.8° in size. The target for the first saccade (referred to as T1) was located 9.7° right of the center (14.6° right of fixation), and the second target (referred to as T2) was located 4.9° left of T1 (9.7° right of fixation). Both targets were white filled circles with a diameter of 0.5° (*Fig. 1A*).

**Figure 1.**
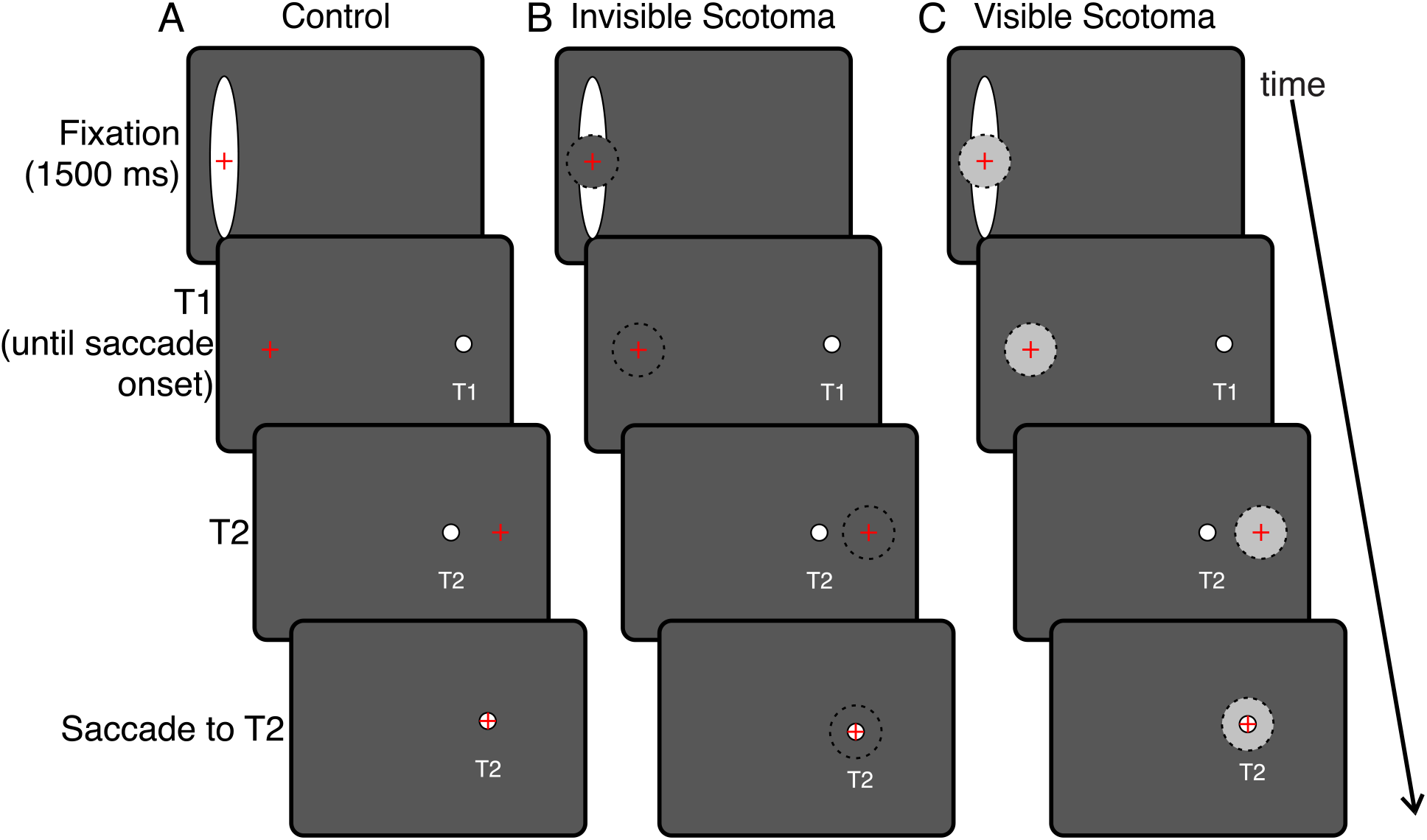
Stimuli and procedure (Experiment 1). The red cross represents the participant’s gaze position. In (B) and (C), the black dotted circle outlines the scotoma. After the second saccade, T2 would be covered by the scotoma and thus not be visible to the participant. The black background of the screen is depicted as dark grey for visibility.

In the scotoma conditions, a black (invisible) or grey (visible) circular central scotoma (3° in diameter) was present. The invisible scotoma was the same colour as the background (i.e. black) and thus not visible (*Fig. 1B*). Its presence was perceived only when occluding stimuli such as the fixation oval. The grey scotoma was visible due to the difference in luminance from the background (*Fig. 1C*). Therefore, it provided information about current eye position to the participant.

Participants took part in three sessions for the adaptation task, completing one of the three conditions (control, invisible scotoma, and visible scotoma) each week in random order. Each session was performed at least one week apart to ensure that there was no retention of adaptation (Alahyane & Pélisson, 2005).

Each session comprised three consecutive blocks. The first block was a pre-adaptation block of 20 trials, in which only T1 was illuminated and extinguished at saccade onset. There was therefore no visual feedback after the first saccade was completed. The second block was the adaptation block, consisting of 180 trials with presentation of both T1 and T2. The last block was the post-adaptation block, which was identical to the pre-adaptation block. The three blocks were run continuously in sequence with no interruption or breaks. In total participants performed 220 trials per session.

In the adaptation block, each trial began with the presentation of the fixation oval which participants were asked to look at (*Fig.1*). After 1500 ms, T1 appeared and participants were instructed to look at it as soon as it appeared. Upon detection of a saccade, T1 was extinguished and T2 was displayed. T2 remained visible for 500 ms. After an inter-trial interval of 500 ms, the fixation oval reappeared and the next trial was initiated.

#### Data Analysis

We collected a total of 7,920 trials from 12 participants. Saccade timing and position were automatically calculated offline using a saccade detection algorithm with a velocity criterion of 15°/s and verified visually. Manual inspection involved removing trials in which saccades were made before the first target appeared, there was a blink during the saccade, the tracker lost eye position, or participants made eye movements not directed toward T1. In total, there were 744 trials removed (9.4% of total trials). We also removed trials in which saccade reaction times were too short (less than 80 ms) or too long (more than 500 ms). There were 127 such trials (1.6% of all trials). Then, we normalized trials in each block by adjusting them by how much the mean saccade start point deviated from fixation point. This was to account for any errors in the calibration process. As mentioned earlier, the accuracy of the eye movement recording was 0.53°. The precision of eye movement recording is much higher (Eyelink reports 0.01° RMS for the Eyelink 1000 Plus). Therefore, while there may be an offset in the eye movement recording, this offset is constant, and precision remains high. We accounted for this offset by normalizing the saccade eye positions for each block.

We removed 28 trials (0.4%) in which participants’ saccades did not begin near the fixation stimulus center (more than 2° away horizontally or vertically) and 2 trials (0.03%) with extremely large saccade amplitude (20° or more). In addition, we removed 128 outlier trials (1.6%) in which the amplitude of the first saccade was more than 2.5 standard deviations away from the mean for each session.

Gain was calculated as the actual saccade amplitude divided by the desired saccade amplitude. The actual saccade amplitude is the difference between horizontal start and end positions of the first saccade. The desired saccade amplitude is the difference between horizontal start position of the first saccade and T1 target position (9.7°). Thus, a gain of one would indicate that the saccade reached T1, and a gain less or greater than one would mean that the participant undershot or overshot the target respectively. We removed 103 gain outlier trials (1.3%) in which gain was more than 2.5 standard deviations away from the mean for each session. In total, there remained 6,788 trials (85.7%).

We calculated the mean gain in the pre-adaptation block and the post-adaptation block for each participant and condition. We determined change in gain for each session as the difference between mean gain in pre-adaptation trials and the mean gain in post-adaptation trials. Also, we calculated the percentage of trials with corrective saccades in the adaptation blocks. Corrective saccades were determined using the following criteria: 1) the start position of the second saccade was less than 1° from the end position of the first saccade, 2) the endpoint of the second saccade was within 5° horizontally of T2, and 3) the saccade had an amplitude greater than 0 and was directed towards T2. Data were analyzed using MATLAB (MATLAB and Statistics Toolbox Release 2018a) and statistical analyses were done with SPSS (SPSS Statistics for Windows, Version 25.0).

### Results

*Figure 2* shows the first saccade endpoints of a typical participant for all three conditions. In all three conditions, there was a shift in saccade endpoints from T1 (dotted line) to T2 (solid line), demonstrating adaptation. Moreover, this was similar across conditions. We also observed that participants tended to undershoot T1 (dotted line) in the pre-adaptation block in all conditions. Interestingly, in the invisible (*Fig. 2B*) and visible (*Fig. 2C*) conditions, the participants’ first saccade endpoints landed so that T2 was occluded by the scotoma (gray region) relatively early in the adaptation block. Nevertheless, adaptation appeared to be the same. These observations are quantified across all participants below.

**Figure 2.**
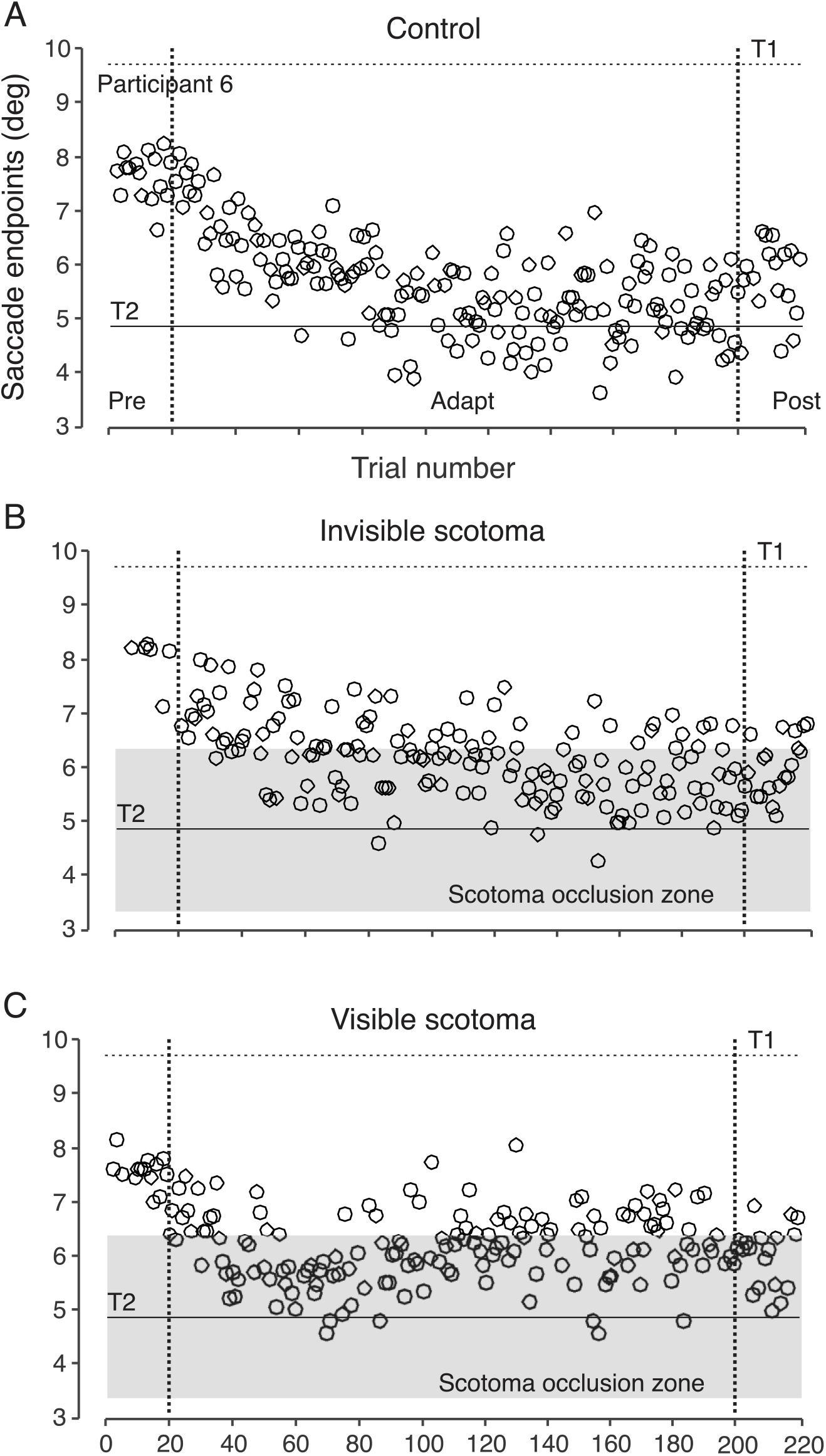
Saccade endpoints of a typical participant (Experiment 1). First saccade endpoints are denoted by black empty circles. Filled in grey is the scotoma occlusion zone, which is 1.5° or less away from T2.

#### Degree of saccadic adaptation

In *Fig. 3A* are depicted the mean gains in the pre- and post-adaptation blocks for each condition. We performed a two-way repeated measures ANOVA with condition (control, invisible, and visible) and block (pre, post) as factors. The change in mean gain for each participant was used, as explained previously. There was a decrease in gain from pre- to post-adaptation blocks in all three conditions, confirmed by a significant main effect for block (*F*(1,11) = 18.7, *p* < 0.001). In addition, we found a significant main effect of condition (*F*(2,22) = 5.99, *p* = 0.008) and a significant interaction effect (*F*(2,22) = 6.84, *p* = 0.005). This indicates that the presence of scotoma and its type affected the amount of adaptation. Post-hoc tests showed that there was a significant decrease in gain in all three conditions. We confirmed that adaptation occurred for the control condition (*Fig. 3*, *left bars*). A paired t-test between mean gain in the pre-adaptation trials (*M* = 0.86, *SD* = 0.05) and post-adaptation trials (*M* = 0.73, *SD* = 0.05) was significant (*t*(11) = 9.49, *p* < 0.001). There was a 14% decrease between the mean gains of pre- and post-adaptation trials, which is about half of the target shift (33% decrease). There was also a significant decrease in the invisible condition (invisible pre *M* = 0.84, invisible post *M* = 0.68, *t*(11) = 13.5, *p* < 0.001) as well as the visible condition (visible pre *M* = 0.81, visible post *M* = 0.71, *t*(11) = 8.29, *p* < 0.001).

**Figure 3.**
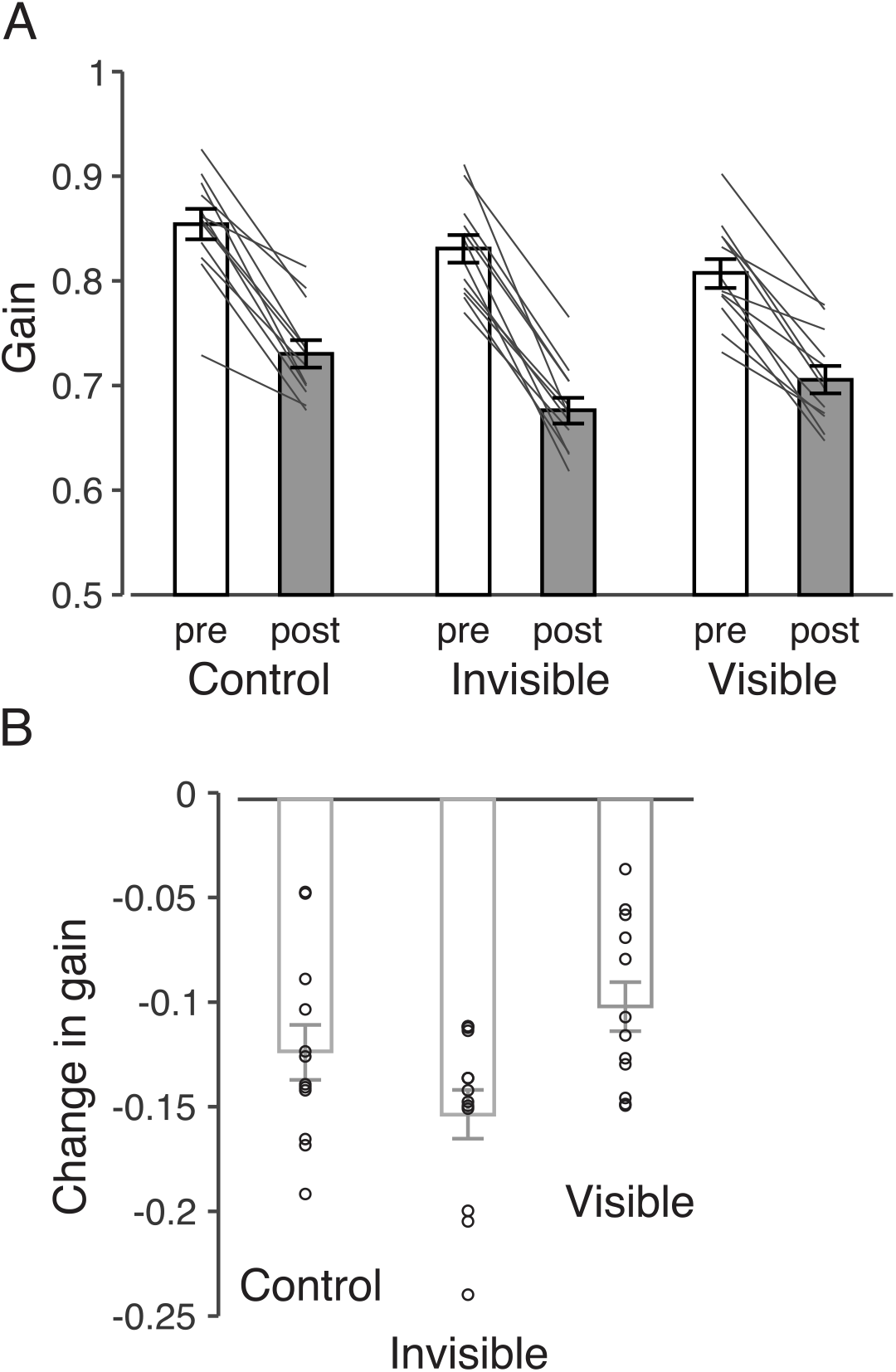
Mean saccade gain by block and condition (Experiment 1). (A) shows the mean gain for the pre-(white bars) and the post-adaptation blocks (gray bars) for each condition as well as individual gains from each participant (thin black lines). (B) shows a bar graph of the change in mean gain between pre-and post-adaptation block for each condition. Open dots represent individual mean gains for each participant.

In addition, we performed two one-way ANOVAs for the pre- and post-adaptation blocks separately. For the pre-adaptation block, the ANOVA was significant (*F*(2,22) = 4.8, *p* = 0.018). Post-hoc t-tests showed significant differences between control (*M* = 0.86) and visible (*M* = 0.81, *t*(11) = 3.32, *p* = 0.007, Bonferroni-Holm familywise error rate) conditions, but no other differences. For the post-adaptation block, the ANOVA was also significant (*F*(2,22) = 8.03, *p* = 0.002). Post-hoc t-tests showed significant differences between control (*M* = 0.73) and invisible (*M* = 0.68, *t(*11) = 3.9, *p* = 0.002) conditions, and between visible (*M* = 0.71) and invisible (*M* = 0.68, *t(*11) = 2.8, *p* = 0.018) conditions. There was no significant difference between control and visible conditions (*p* = 0.16). In summary, it appears that participants had smaller gains in the pre-adaptation block of the visible condition, possibly due to visual feedback of eye position. This was not the case in the post-adaptation block.

To compare difference in adaptation, we compared change in gain between pre- and post-adaptation blocks for the three conditions (*Fig. 3B*). First, we confirmed that there was significant change in gain through one-sample t-tests. All three conditions showed changes in gain that were significantly different from 0 (all *p* < 0.001). A one-way repeated-measures ANOVA on gain change with condition as a factor was significant (*F*(2,22) = 6.83, *p* = 0.005). Post-hoc testing revealed that gain reduced to a greater degree for the invisible (*M* = −0.15) compared to the visible (*M* = −0.12, *t(*11) = 3.3, *p* = 0.007) condition, but there were no differences between invisible and control (*M* = −0.12, *p* = 0.043, Holm-Bonferroni family-wise error rate = 0.025), nor between control and visible (*p* = 0.13). In summary, adaptation was largest for the invisible condition and smallest for the visible condition, with no significant differences from control, whose gain was in between the two.

#### Occlusion of the 2^nd^ target by the scotoma

As shown in *Fig. 2A* and *B*, around midway in the adaptation block, many saccade endpoints landed within the scotoma occlusion zone. In other words, for any saccade endpoint that landed at 6.35° or less, the scotoma occluded T2. It appears that this did not impact adaptation, however. We calculated the percentage of saccade endpoints that landed within this zone for all participants. The amount of occlusion was quite substantial, ranging from 47% to 96% for the invisible condition (*M* = 84%, *SD* = 15.7%) and 34% to 98% for the visible condition (*M* = 73%, *SD* = 25.5%). We compared each participant’s amount of gain change and the percent of occlusion for each condition to investigate whether increased occlusion led to decreased adaptation. As expected, given that the invisible condition showed more adaptation with more occlusion, we did not find a significant relationship for either condition (*p* > 0.05).

In order to determine when the target was occluded relative to the ongoing saccade during the adaptation block, we first calculated when T2 appeared relative to saccade onset. On average, T2 appeared 37.1 ms (*SD* = 3.3 ms, across participants) after saccade onset in the control condition, 38.8 ms (*SD* = 3.6 ms) in the invisible condition and 37 ms (*SD* = 3.7 ms) in the visible condition. We confirmed that there were no significant differences across conditions through a repeated-measures ANOVA (*p* > 0.05). With respect to saccade peak velocity, T2 appeared very slightly ahead of peak velocity time of the saccade, appearing on average 2.3 ms before (*SD* = 4.8 ms) for the control condition, 3 ms (*SD* = 5.3 ms) for the invisible condition and 2.6 ms (*SD* = 4.3 ms) for the visible condition. Again, there were no differences across conditions (*p* > 0.05). Next, we calculated the duration of T2 visibility, which was the time between when T2 appeared and when it was occluded by the scotoma (during the saccade). In other words, the latter is the point during which the saccade was at the T2 position minus 1.5°. On average, before being occluded by the scotoma, T2 appeared for 18.7 ms (*SD* = 8.6 ms) in the control condition, 17.6 ms (*SD* = 7.2 ms) in the invisible condition, and 16.5 ms (*SD* = 6.9 ms) in the visible condition. As before, there were no differences across conditions (*p* > 0.05). Note that these calculations were made for all saccades. To summarize, T2 appeared mostly during the deceleration phase of the saccade, from just before peak velocity. As described above, only a certain percentage of these saccades had amplitudes for which T2 remained occluded even at the end of the saccade. Other saccades had larger amplitudes, so that by the end of the saccade T2 was no longer occluded. In short, viewing T2 for approximately 15 ms during the later stages of the saccade was sufficient to drive adaptation, as previously shown (Panouillères et al., 2013).

#### Corrective saccades

We investigated whether there was a relationship between the number of corrective saccades performed and the amount of adaptation. We observed that across all conditions, half (6) the participants made no corrective saccades. Two participants made minimal corrective saccades in one of the conditions (participant 5, control condition, 5%; participant 6, invisible condition, 8%). For the four remaining participants, 69% of all trials comprised corrective saccades (*SD* = 23%) for the control condition, 41% (*SD* = 31%) for the invisible condition, and 41% (*SD* = 41%) for the visible condition.

There were no significant differences overall across the conditions (*F*(2, 22) = 3.29, *p* = 0.06). Moreover, there was no significant relationship between mean change in gain and the percentage of corrective saccades in any of the 3 conditions (*p* > 0.05). These results show that corrective saccades did not play a role in adaptation.

#### Experiment 1 Summary

Participants showed backward saccadic adaptation in all three conditions. Overall, the amount of adaptation was similar across the three conditions, even with some feedback of T2 during the later stages of saccades. These results show that occlusion of central vision does not affect adaptation. However, a concern with the backward adaptation paradigm is that fatigue may have caused a large proportion of saccade gain decrease. Although extraocular muscles tend to be relatively resistant to fatigue (Fuchs & Binder, 1983; Saito, 1992), it has been shown in both humans (De Gennaro et al., 2000, 2001; Rowland et al., 2005; Sprenger et al., 2005) and monkeys (Straube et al., 1997; Straube et al., 1997) that fatigue can affect saccade metrics. Therefore, the amounts of adaptation during the scotoma conditions might be related to fatigue rather than adaptation per se. In addition, our results for backward adaptation may not be generalizable to forward adaptation. Backward and forward adaptation are likely based on different mechanisms (Catz et al., 2008; Ethier et al., 2008; Golla et al., 2008; Hernandez et al., 2008; Pélisson et al., 2010; Semmlow et al., 1989). Therefore, we cannot make conclusions about forward adaptation based on results from this first experiment.

In order to address these outstanding issues, we performed a second experiment in which forward adaptation was tested. In this experiment, we used two differently sized invisible central scotomas. By varying the diameter of scotoma, we could investigate how the eccentricity of the viewed peripheral T2 affects adaptation. A larger scotoma results in larger eccentricities of T2 relative to the fovea that are not occluded.

## Experiment 2

### Methods

This experiment was almost identical to Exp. 1, with a few changes that are outlined below.

#### Participants

Thirteen participants took part in this study (four male, age range: 19-42, *M* = 24.38, *SD* = 5.84, including the author AK). All participants had normal or corrected-to-normal vision, no known neurological impairments, and gave written informed consent to participate in the experiment.

#### Procedure

Stimuli were presented with custom code using MATLAB (The Math Works Inc., Natick, MA) with Psychtoolbox (Brainard & Vision, 1997; Pelli & Vision, 1997) and EyeLink toolboxes (Cornelissen et al., 2002). The procedure was the same as described in Exp. 1. There were three conditions in total: a control condition with no scotoma, a 2° diameter scotoma condition, and a 4° diameter scotoma condition. Aside from the presence of a scotoma and its size, all conditions were identical.

Stimuli used are shown in *Figure 4*. The white fixation oval was identical to that in Exp. 1. The target for the first saccade (referred to as T1) was located 5° right of the center (10° right of fixation), and the second target (referred to as T2) was located 5° right of T1 (15° right of fixation). Both targets were white filled circles with a diameter of 0.5°.

**Figure 4.**
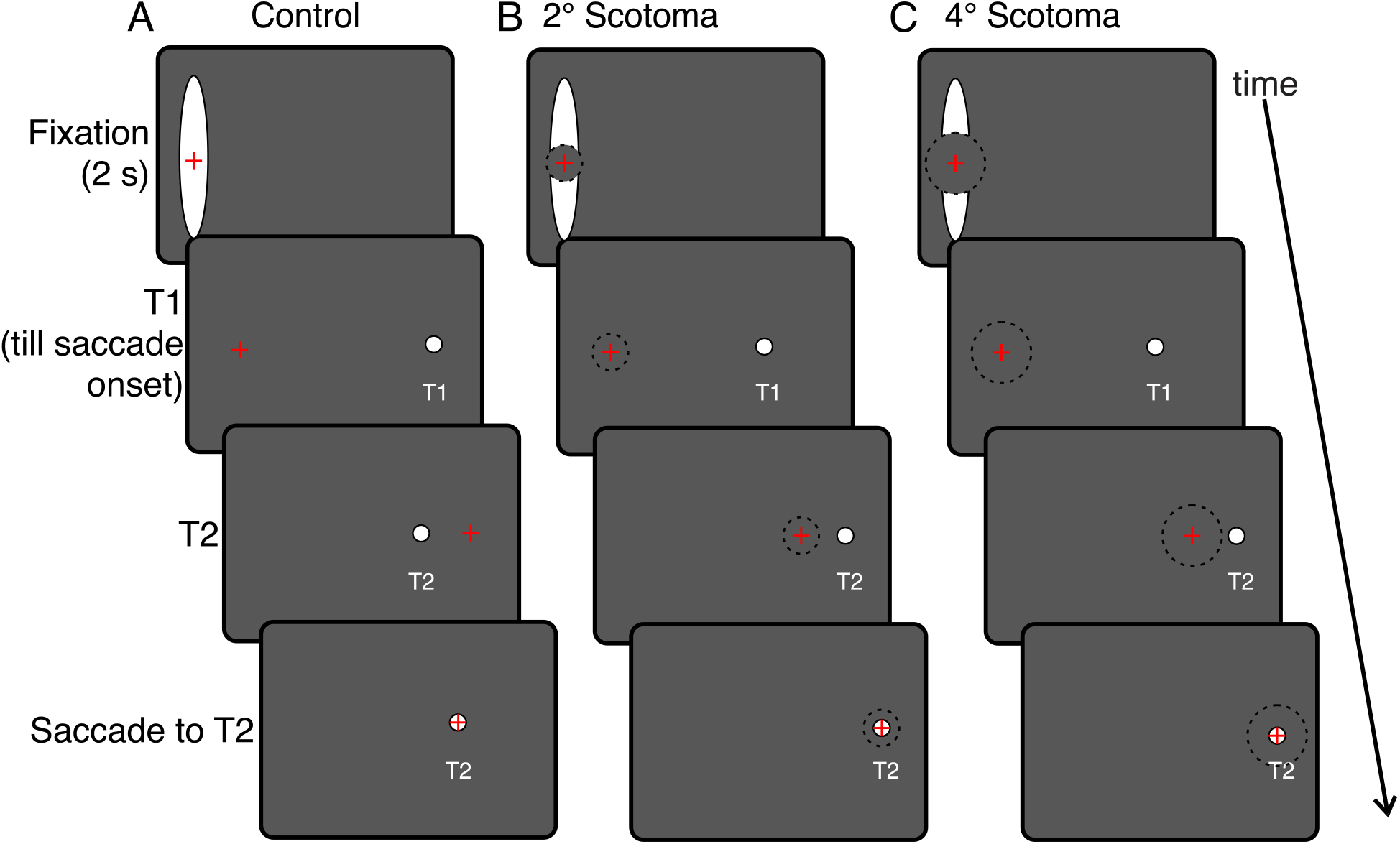
Stimuli and procedure (Experiment 2). The red cross represents the participant’s gaze position. In (B) and (C), the black dotted circle outlines the scotoma. The black background of the screen is depicted as dark grey for visibility.

In the scotoma conditions (*Fig. 4B* & *C*), a black circular central scotoma (2° or 4°) was present. It was the same colour as the background (i.e. black), and so, was invisible. Its presence was perceived only when occluding stimuli such as the fixation oval.

Participants took part in three sessions completing each at least a week apart in random order. Each session comprised three blocks. The first block was a pre-adaptation block, comprising 25 trials. In this block, only T1 was illuminated and extinguished at saccade onset. The second block was the adaptation block, comprising 200 trials with the presentation of both T1 and T2. The last block was the post-adaptation block, which was identical to the pre-adaptation block. In total participants performed 250 trials per session, which took 12 to 15 minutes. We increased the number of trials from the first experiment as forward adaptation typically takes longer than backward adaptation (Lévy-Bencheton et al., 2016).

Each trial began with the presentation of the fixation oval at which participants were asked to look. After 2000 ms, T1 appeared and participants were instructed to look at it as soon as it appeared. When a saccade was detected, T1 was extinguished and T2 was displayed. On average, T2 appeared 2.25 ms after the time of peak saccade velocity (beginning of the deceleration phase). T2 remained visible for 400 ms. For the two scotoma conditions, T2 was also presented for 400 ms, although it was not visible after the corrective saccade as it was covered by the scotoma. After an inter-trial interval of 400 ms, the fixation oval re-appeared and the next trial was initiated.

#### Data Analysis

The same parameters and analysis methodology were used as in Exp. 1. We collected a total of 9,747 trials from 13 participants. Trials were removed in which 1) saccades were made before the first target appeared, there was a blink during the saccade, the tracker lost eye position, or participants made eye movements not directed toward T1 (1,247 trials, 12.8% of total trials), 2) saccade reaction time was not between 80 and 500 ms (214 trials, 0.2%), 3) participant’s normalized saccades did not begin near fixation stimulus center (122 trials, 0.13%), 4) first saccade amplitude was more than 2.5 standard deviations away from the mean for each session (144 trials, 0.15%), and 5) gain was more than 2.5 standard deviations away from the mean for each session (107 trials, 0.11%). In total, there remained 7,913 trials (81.2%).

### Results

*Figure 5* depicts saccade endpoints for a typical participant in all three conditions, showing similar amounts of adaptation across the three conditions.

**Figure 5.**
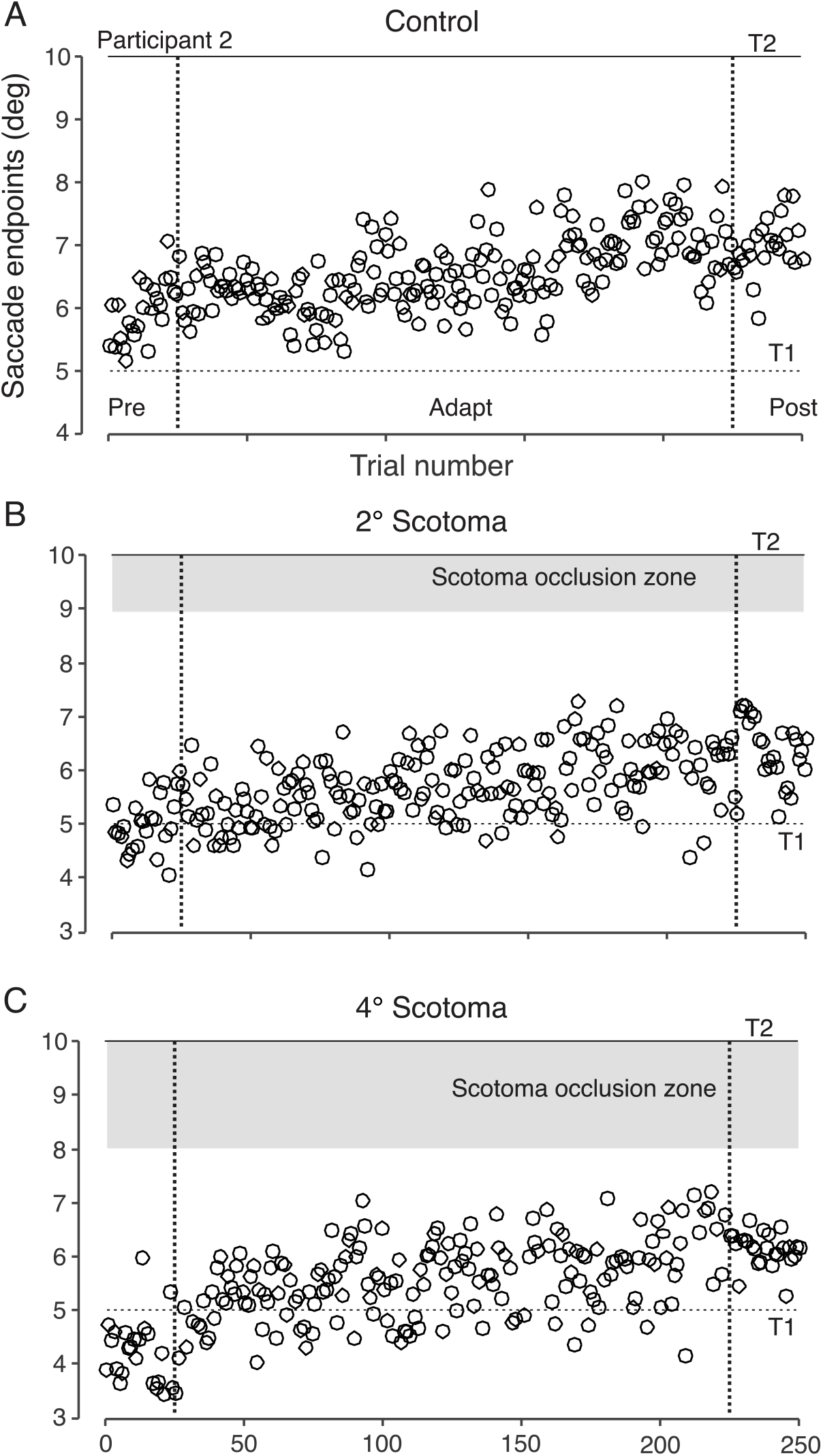
Saccade endpoints of a typical participant (Experiment 2). First saccade endpoints are denoted by black empty circles. Filled in grey is the scotoma occlusion zone, which is 1° or less away from T2 in (B) and 2° or less away from T2 in (C). Note that in both (B) and (C) there were no endpoints in the occlusion zone, in contrast to Experiment 1.

#### Saccadic adaptation

In *Fig. 6A* can be seen the mean gains for the pre- (white bars) and post-adaptation (gray filled bars) blocks for each condition as well as individual gains (thin black lines). We observed that participants were less consistent in demonstrating forward adaptation compared to backward adaptation in Exp. 1. Some individual participants even showed a decrease in gain. We performed a two-way repeated measures ANOVA on change in mean gain with condition (control, 2°, and 4°) and block (pre, post) as factors. The ANOVA revealed a significant main effect of condition (*F*(2,24) = 3.8, *p* = 0.037) and a significant main effect of block (*F*(1,12) = 22.7, *p* < 0.001), but no interaction effect (*p* > 0.05). These results suggest that there was a significant increase in gain for all three conditions (mean pre gain = 0.99, mean post gain = 1.06). Bonferroni-corrected pairwise comparisons also revealed that the overall gain (collapsed across pre and post) was largest for the 2° scotoma condition (*M* = 1.05), which was significantly different from the 4° scotoma condition (*M* = 1, *p* = 0.01), but not from control (*M* = 1.02).

**Figure 6.**
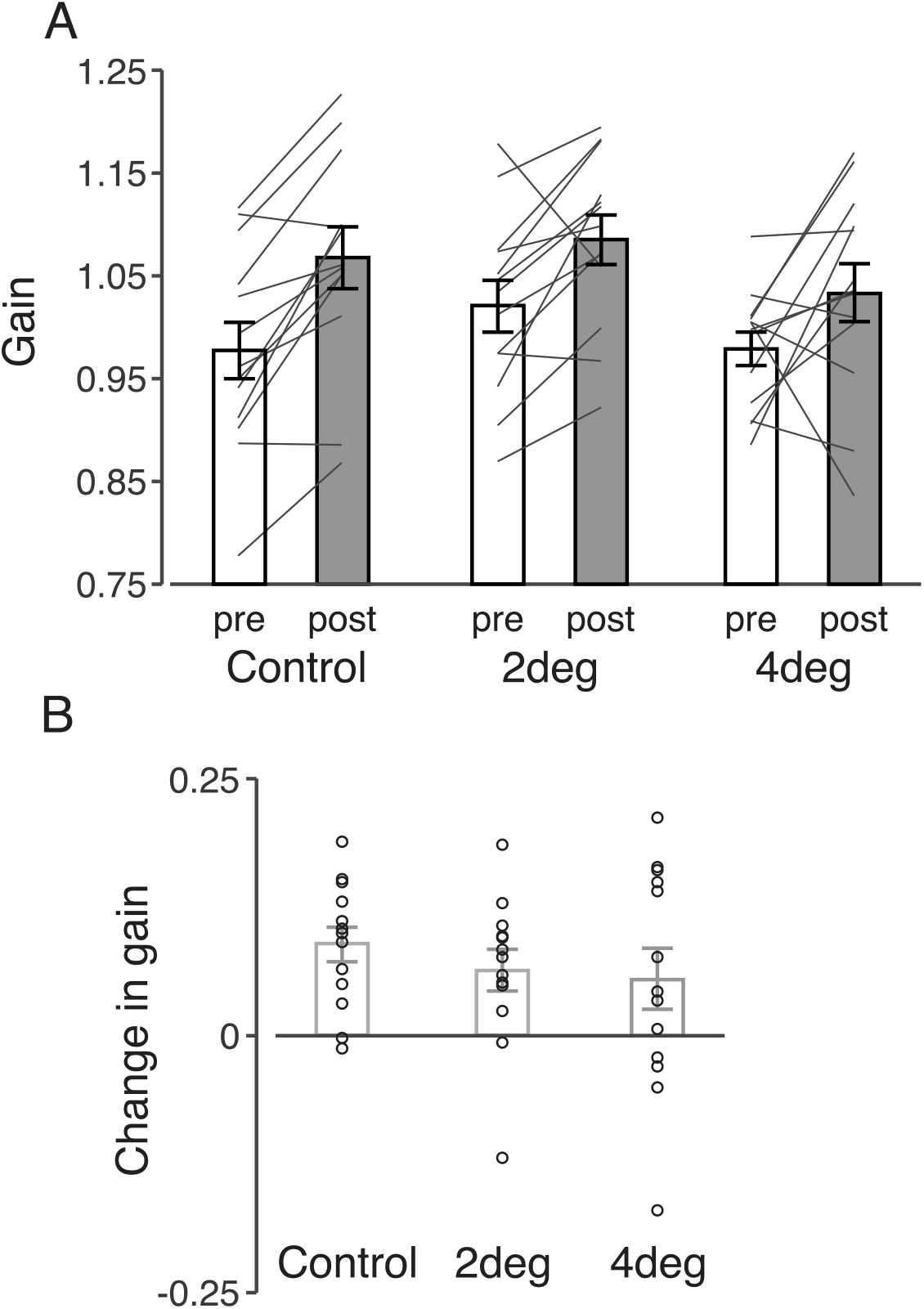
Mean saccade gain by block and condition (Experiment 2). Data are presented in the same manner as in Fig. 3.

In *Fig. 6B* can be seen the mean (bars) and individual (dots) change in mean gain for the three conditions. Interestingly, we observed with one-sample t-tests that the change in mean gain was not significantly different from 0 for the 4° scotoma condition (*t(*12) = 1.8, *p* = 0.09), while it was significantly different for the other two conditions (control, *t(*12) = 5.3, *p* < 0.001; 2°, *t(*12) = 3.2, *p* = 0.008). Nevertheless, the change in mean gain was not significantly different across conditions, as shown by a one-way ANOVA (*F*(2,24) = 0.6, *p* = 0.5, control *M* = 0.089, 2° *M* = 0.064, 4° *M* = 0.055).

#### Scotoma occlusion

Unlike Exp. 1, very few saccade endpoints landed such that the scotoma occluded T2. For the 2° scotoma condition, 12 out of 13 participants had no saccades landing within this zone during the adaptation block and one participant had only 0.5% of saccades. For the 4° scotoma condition, only 3 out of 13 participants had saccades landing within the zone (*M* = 2.12%, *SD* = 0.63%).

#### Corrective saccades

Like in Exp. 1, we compared the proportion of corrective saccades with the change in gain across participants. Overall, there was a wide range in the percentage of adaptation trials with corrective saccades overall (*M* = 76.3%, *SD* = 34.8%), ranging from 100% to 1.4%, but a few participants were responsible for most of the variability. In particular, participant 6 made very few corrective saccades (<14% in the three conditions) as did participant 13 (<15%). The remaining 11 participants made corrective saccades in most trials. This contrasts with Exp. 1 in which most participants did not make corrective saccades.

However, similar to Exp. 1, there were no significant differences across conditions (control, *M* = 76.3%, *SD* = 38.6%; 2°, *M* = 77.2%, *SD* = 35.6%; 4°, *M* = 75.2%, *SD* = 32.7%; *F*(2.24) = 0.098, *p* = 0.9). Moreover, there was no significant relationship between mean change in gain and the percentage of corrective saccades in any of the 3 conditions (*p* > 0.05). Thus, corrective saccades did not play a role in adaptation.

#### Experiment 2 summary

Presence of the invisible scotoma (2° and 4°) resulted in similar amounts of adaptation compared to control, confirming results from Exp. 1 that occlusion of central vision does not affect saccadic adaptation. Unlike Exp. 1, we observed that there was almost no occlusion of T2 after the first saccade. This was because most saccades undershot T1, as is the general tendency for saccades (Becker & Fuchs, 1969; de Bie et al., 1987; Deubel et al., 1982; Kapoula, 1985).

### Discussion

We measured the extent to which saccadic adaptation occurred in the presence of artificial central scotomas of different visibilities and sizes in both forward and backward paradigms. We observed similar amounts of adaptation between scotoma and control conditions. In the backward adaptation paradigm, we observed adaptation even when the scotoma occluded the shifted target most of the time at the end of the saccade. Adaptation took place in response to the shifted target during the later stages of the ongoing saccade. In the forward adaptation paradigm, we also observed similar amounts of adaptation compared to control for two differently sized invisible scotomas. We conclude that saccadic adaptation occurs equally well in the presence of a central scotoma and that peripheral peri- and post-saccadic visual feedback of the shifted target location are sufficient to drive adaptation.

In the backward adaptation paradigm, the central scotoma often occluded T2 at the end of the first saccade, impeding adaptation. However, we observed that adaptation occurred equally well. There was similar adaptation across the three conditions, confirming previous findings (Panouillères et al., 2016; Panouillères et al., 2013). Specifically, it has been shown that intra-saccadic visual feedback received mid-flight during a saccade can cause adaptation (Panouillères et al., 2016; Panouillères et al., 2013). The effect of visual feedback timing was tested by comparing an intra-saccadic condition, in which the shifted target was displayed only during the saccade, and a post-saccadic condition, in which the shifted target was displayed after the saccade (Panouillères et al., 2013). The two conditions produced equal amounts of adaptation, for both backward and forward target shifts. In an additional experiment using backward target shifts, even displaying the target for 10 ms or 2 ms durations was sufficient to cause adaptation in the same manner as post-saccadic presentation, but only during the deceleration phase of the saccade and not acceleration phase or at peak velocity (Panouillères et al., 2016). To summarize, it appears that even intra-saccadic *peripheral* presentation of T2 is sufficient for adaptation and post-saccadic foveal or peri-foveal information does not increase the amount of adaptation.

While there was a consistent decrease in gain across all three conditions of the backward paradigm, there was less consistent gain change in the forward paradigm, with some participants even showing gain decrease. This is consistent with previous findings which show that forward adaptation does not always result in gain increase, particularly for target shifts of less than 50% of target eccentricity (Robinson et al., 2003). Other studies have also demonstrated that a larger number of trials are needed to elicit gain increase (equal in magnitude to gain decrease in backward adaptation) in forward adaptation (Deubel et al., 1986; Miller et al., 1981; Straube et al., 1997). It has been proposed that forward and backward adaptation are based on different mechanisms in the brain (Ethier et al., 2008; Hernandez et al., 2008; Pélisson et al., 2010; Semmlow et al., 1989) which would likely lead to different behavioral patterns.

We also investigated differences in the number of corrective saccades across conditions and if this was related to the amount of adaptation. In both cases, we found no differences and no relationship, confirming that corrective saccades did not play a role in adaptation. Specifically, it appears that making a corrective saccade to T2 after the first saccade to T1 did not play a role in determining the error for which the saccade must compensate. In addition, the lack of difference across all conditions demonstrates that the scotoma did not influence corrective saccades. As for how corrective saccades would influence adaptation, most previous studies show they are insignificant: removing corrective saccades had almost no effect on saccade gain and changing the direction of corrective saccades had no influence either (Noto & Robinson, 2001; Wallman & Fuchs, 1998).

For both backward and forward adaptation, we observed that foveal feedback of T2 was not important. In the backward paradigm, neither the visible nor the invisible scotoma condition was different from control in the amount of adaptation. In the forward paradigm, there was no significant difference across conditions in the amount of adaptation. While it is well established that foveal feedback of the shifted target is unnecessary for adaptation, the nature of the error signal that drives adaptation is still unresolved. It was once proposed that visual retinal error (how far off the fovea is from the target post-saccade) drove adaptation (Noto & Robinson, 2001; Wallman & Fuchs, 1998). Recent studies suggest that adaptation is caused not by retinal error per se, but by the difference between the retinal image (post-saccade) and the predicted image (pre-saccade), also referred to as the visual comparison model (Bahcall & Kowler, 2000). Retinal error is not an adequate error signal in a real world scenario, such as scanning scenery in nature, in which it would be difficult to determine retinal error because of the numerous visual objects that can take on a variety of shapes (Bahcall & Kowler, 2000). Bahcall and Kowler (2000) also demonstrated saccadic adaptation during a task in which participants were instructed to saccade partway to a target (75% of the distance from initial fixation point). In this case, the retinal error would always be positive. However, the target was shifted backwards, and this resulted in gain decrease rather than increase as adaptation driven by retinal error would suggest. The visual comparison model is analogous to the more general sensory prediction error (SPE) hypothesis proposed by some to drive adaptation, defined as the discrepancy between predicted and actual sensory signals (Herman et al., 2013). There is evidence for sensory prediction errors in the visuomotor system (Mazzoni & Krakauer, 2006; Shadmehr et al., 2010; Tseng et al., 2007) as well as visual perception (Alink et al., 2010; Den Ouden et al., 2012; Meyer & Olson, 2011).

We observed no significant difference in the amount of mean gain change between control and scotoma conditions, regardless of the size of the scotoma, albeit we tested using relatively small diameters. These results support the idea that saccadic adaptation could be possible in patients with AMD as a means of training. We speculate that adaptation could be used to adapt eye position such that the desired target lands on the PRL after the first saccade by training for numerous trials. We also found similar adaptation with both visible and invisible central scotomas, suggesting that adaptation could occur even when patients are unaware of the presence of the scotoma itself (Fletcher et al., 2012). It has been suggested however that scotoma awareness is a possible tool for rehabilitation for patients with central vision loss (Scheiman et al., 2007; Walsh & Liu, 2014). This could be achieved using a gaze-contingent visible scotoma with the shape of the patient’s scotoma but slightly larger and may aid patients in reinforcing adaptation to make optimal use of their peripheral vision (Barraza-Bernal et al., 2017; Walsh & Liu, 2014). Generally, the effects of saccadic adaptation have been shown to remain for a short duration, typically under a week (Alahyane & Pélisson, 2005). But, this could be because saccades which direct the object of interest outside the fovea are not optimal for healthy participants. If AMD patients are trained so that saccades direct targets to the PRL, it could be beneficial to them, and therefore the effects of adaptation might be reinforced and better maintained.

In conclusion, we showed that both backward and forward adaptation occurred equally well in the presence of an artificial gaze-contingent central scotoma as without. We propose using saccadic adaptation as a means of training saccade re-referencing for people with central vision loss.

## Acknowledgements

AZK was funded by the Canada Research Chair program and the National Sciences and Engineering Research Council of Canada

